# Uncovering modeling features of viral replication dynamics from high-throughput single-cell virology experiments

**DOI:** 10.1101/2020.07.09.195925

**Authors:** Ashley I. Teufel, Wu Liu, Jeremy A. Draghi, Craig E. Cameron, Claus O. Wilke

## Abstract

Viruses experience selective pressure on the timing and order of events during infection to maximize the number of viable offspring they produce. Additionally, they may experience variability in cellular environments encountered, as individual eukaryotic cells can display variation in gene expression among cells. This leads to a dynamic phenotypic landscape that viruses must face to produce offspring. To examine replication dynamics displayed by viruses faced with this variable landscape, we have developed a method for fitting a stochastic mechanistic model of viral infection to growth data from high-throughput single-cell poliovirus infection experiments. The model’s mechanistic parameters provide estimates of several aspects associated with the virus’s intracellular dynamics. We examine distributions of parameter estimates and assess their variability to gain insight into the root causes of variability in viral growth dynamics. We also fit our model to experiments performed under various drug treatments and examine which parameters differ under these conditions. We find that parameters associated with translation and early stage viral replication processes are essential for the model to capture experimentally observed dynamics. In aggregate, our results suggest that differences in viral growth data generated under different treatments can largely be captured by steps that occur early in the replication process.

**Author Summary:** Understanding the intercellular processes associated with virus replication is essential for controlling viral diseases. Single-cell infection experiments with poliovirus have shown that viral growth differs among individual cells in terms of both the total amount of virus produced and the rate of viral production. To better understand the source of this variation we here develop a modeling protocol that simulates viral growth and we then fit our model to data from high-throughput single-cell experiments. Our modeling approach is based on generating time course simulations of populations of single-cell infections. We aggregate metrics from our simulated populations such that we can describe our simulated viral growth in terms of several distributions. Using the corresponding distributions calculated from experimental data we minimize the difference between our simulated and experimental data to estimate parameters of our simulation model. Each parameter corresponds to a specific aspect of the intercellular viral replication process. By examining our estimates of these parameters, we find that steps that occur early in the viral replication process are essential for our model to capture the variability observed in experimental data.

## Introduction

Populations of RNA viruses are composed of distinct genetic variants that compete within host cells [1–4]. The host cells themselves display a significant amount of variability in gene expression [5–7], causing the competition of RNA variants to occur within a dynamic system. Genetic variability in viral populations coupled with variability in host cell conditions renders the process of viral replication highly stochastic in nature. The stochasticity of the replication process has been well documented [8, 9] and has been observed in high-throughput single-cell poliovirus (PV) infections experiments [10–12]. By measuring viral replication across the infection process under various conditions, these types of high-throughput experiments can gather a wealth of metrics about the timing and progression of infection. These metrics can be examined and compared across experimental conditions to enable an in-depth look at the process of viral replication.

Understanding the molecular events accompanying virus replication is essential for the control of all viral diseases [13]. Knowledge of the replication processes and mutation rates can yield insight into managing drug resistance, immune escape, vaccination, pathogenesis, and the emergence of new diseases [14]. The data generated from high-throughput single-cell experiments provide a rich source of information for measuring changes in the replication process under various treatments, but they lack the capacity to describe the underlying intracellular dynamics leading to the observed differences [15]. The ability to quantify how intracellular dynamics differ across treatments can lead to insight about how specific treatments impact the intracellular replication process differently. For example, if a drug treatment increases the length of time it takes before an infected cell lyses, this may have downstream effects on other aspects of the viral replication process. However, measuring how a treatment specifically impacts intracelluar dynamics in-vivo is a challenging task [16].

To examine the intracellular dynamics of viral infection, we have developed a method for fitting population-level viral growth data from high-throughout single-cell experiments to a mechanistic model of poliovirus replication. Our method simulates hundreds of individual viral growth trajectories in single cells to generate population-level data. We estimate the parameters in the mechanistic model by minimizing the difference between distributions of four metrics that describe the dynamics observed in populations of infected cells. We then examine the distributions of mechanistic parameter estimates to assess which of these parameters are informative about intercelluar dynamics. We find that parameters associated with translation and early stage replication processes are essential for describing viral growth. To further assess which aspects of our model are important for fitting experimentally generated viral growth data, we use our model fitting method to fit growth data from experiments under three different drug treatments. These drug treatments include a 3C protease inhibitor (rupintrivir), a polymerase inhibitor (2’-C-meA), and a Hsp90 inhibitor (ganetespib). We again find that the same parameters appear to control our model’s ability to capture realistic growth dynamics. Our results suggest that rates of some specific aspects of the viral replication process can be estimated using this modeling approach, and that these rates can inform us about aspects of the replication process that are critical for generating variation in population-level viral growth data.

## Results

### Characterizing population-level infection dynamics from single-cell data

We make use of data sets generated by [12] of poliovirus (PV) encoding GFP (GFP-PV), infecting individual HeLa S3 cells. GFP-PV was created by inserting GFP-coding sequence between the capsid-coding P1 region of the PV genome and the non-structural protein-coding P2 region. Experiments using this GFP-PV have been performed using a microfluidic device capable of capturing thousands of potential infection events in single cells using florescence detection. Changes in florescence are monitored over time with time-lapse microscopy. These time-lapse data are analyzed by fitting either a sigmoidal function (modeling no lysis, Fig. 1A) or a double sigmoidal function (modeling lysis, Fig. 1B) to each infection event [17]. For each infection, one or the other of these two fitted functions is selected as the best fit, based on a model-fit criterion [17]. This fitting procedure allows us to quantify the distributions of parameters: the slope of the first sigmoidal function at the midpoint (slope), the maximum amount of GFP intensity reached (maximum), the midpoint at which half of the maximum GFP intensity is reached (midpoint), and the lysis time (lysis) (Fig. 1A, B). By aggregating these parameters over many individual infections, we can estimate the distributions of each of these parameters (Fig. 1C–F). These distributions can then be used to compare population-level infection dynamics generated under different experimental conditions.

**Figure 1:**
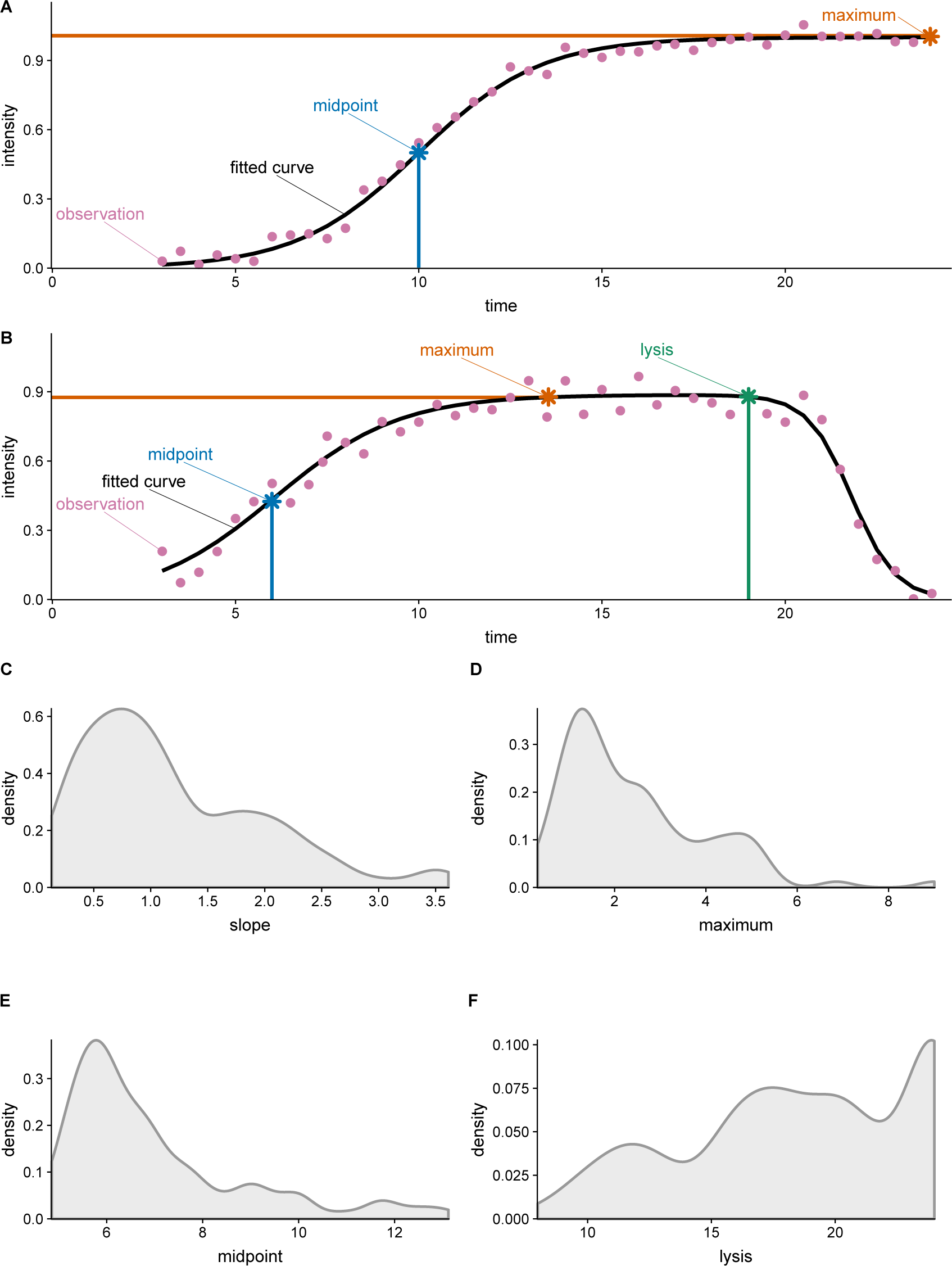
Sigmoidal functions fit to experimental data and distributions of fitted parameters when the sigmoidal functions are fit to a population of infection events generated by [12] without drug treatment. (A) Example of sigmoidal function fit to experimentally data. A sigmodal curve (black line) is fitted to intensity observations from a single infection event (purple dots), allowing for the estimation of the midpoint (blue) and the maximum intensity value (orange). (B) Example of double-sigmoidal function fit to a single infection event where lysis is observed. A double sigmodal curve (black line) is fitted to intensity observations from a single infection event (purple dots), allowing for the estimation of the midpoint (blue), the maximum intensity value (orange), and the time point of lysis (green). (C) Distribution of slopes calculated at the midpoint. The slope is related to the viral replication rate. (D) Distribution of the maximum intensity. (E) Distribution of the amount of time until the half of the maximum intensity is reached. (F) Distribution of the length of infection time.

### Model of intercelluar replication

Here, we have modified a previously published model of PV replication [18] so it can be fitted to the distributions of parameters (slope, max, midpoint, and lysis) collected from the single-cell experiments, and we show that we can use this approach to characterize variation in PV infections among individual cells and under different experimental conditions. [18] previously introduced a stochastic model that simulates individual reactions and tracks abundances of PV molecular species within a cell. The model consists of eight distinct steps (Figure 2A). The full mathematical description of this model is presented in [18] and is summarized in Table 1. Here, we briefly describe the biological meaning of each step.

**Table 1:**
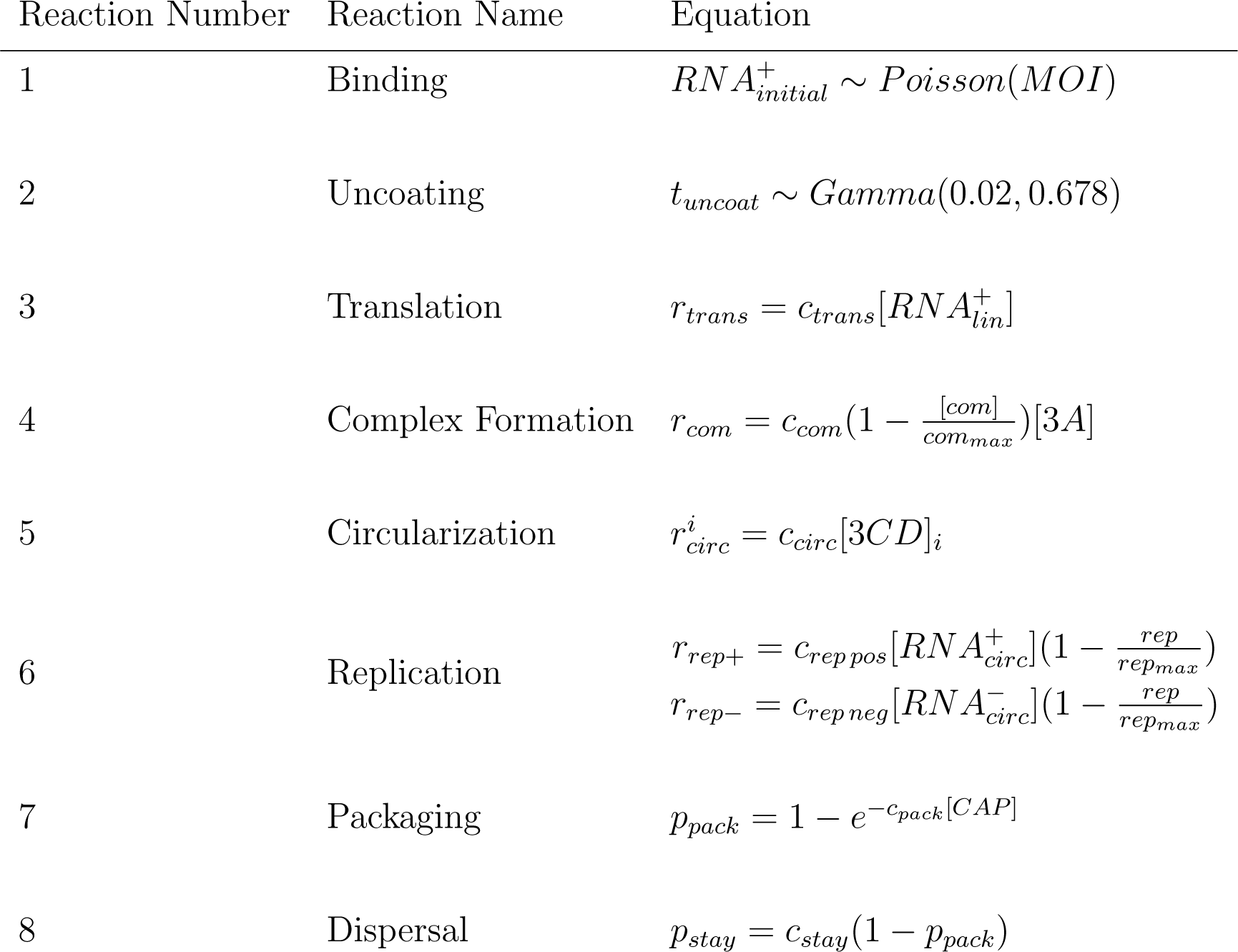
The reactions that describe the replication cycle of PV. Numbered steps correspond to individually modeled reactions as described in Figure 2. See [18] for a full mathematical description of the model.

**Figure 2:**
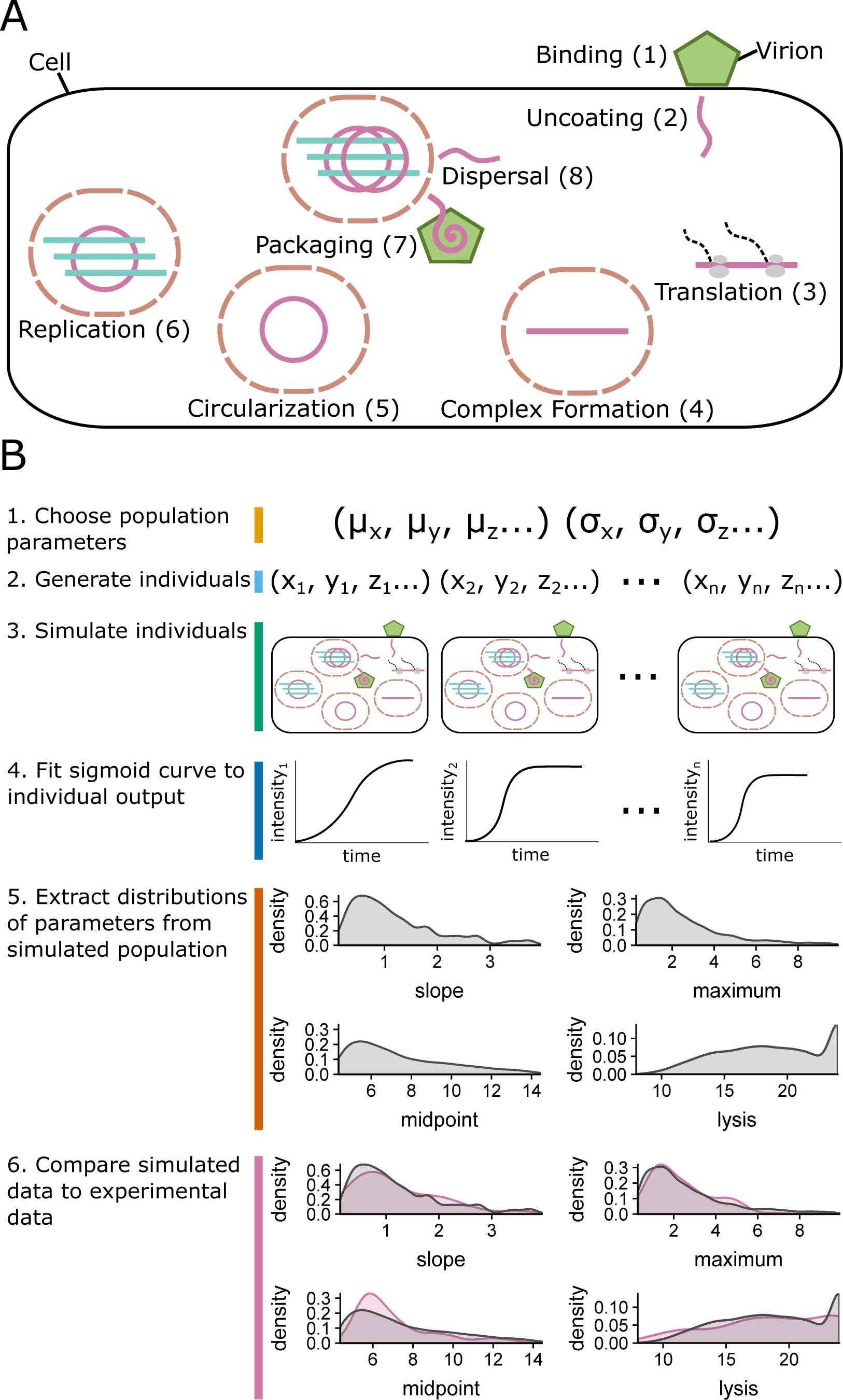
Illustration of PV replication cycle and parameter estimation procedure. (A) The replication cycle of PV as represented in the model of [18]. This figure is adapted from that work. Numbered steps correspond to individually modeled reactions given in Table 1. (B) Computational procedure to compare the output of the mechanistic model of a single PV infection to the experimental data obtained from thousands of single-cell infections.

First, infecting virions bind to and infect a cell (Fig. 2A, step 1). The binding process is based on a population’s average multiplicity of infection (MOI) and the number of infecting virions is drawn from a Poisson distribution with a mean equal to the MOI. Uncoating occurs after a gamma-distributed delay occurs [19], resulting in positive-sense genomes being introduced into the cell (Fig. 2A, step 2). Once positive-sense genomes are present translation begins (Fig. 2A, step 3), dependent on a parameter that describes translation (*c*_*trans*_) and the amount of RNA genomes present.

After a sufficient amount of protein products are present, the positive sense strands form replication complexes (Fig. 2A, step 4). Complex formation is a function of a parameter that describes complex formation (*c*_*com*_), the amount of the protein product bounded by the maximum amount of possible complexes that could be formed (*com*_*max*_), and the amount of a protein product (3*A*) that facilitates this complex formation. The genomes can then circularize to become competent for replication (Fig. 2A, step 5), and this process is modeled as a function of a parameter that describes circularization (*c*_*circ*_) and the amount of a protein needed for the genome to circularize within the complex (3*CD*).

Replication of negative-sense then positive-sense genomes occurs (Fig. 2A, step 6) after a positive-sense genome has circularized. The rates of production of newly synthesized negative and positive sense genomes are a function of parameters that describe the rate of positive and negative strand synthesis (*c*_*rep-*_ and *c*_*rep*_+, respectively). This replication process depends on the amount of positive and negative sense circularized genomes, and is bounded by the maximum possible number of replication cycles (*rep*_*max*_). The *rep*_*max*_ parameter reflects resource limitations within the cell. The equation that describes the rate of positive strand replication accounts for the rate at which replication is initiated on positive-sense templates, leading to the productions of negative-sense strands. Similarly, the equation that describes the rate of negative strand replication accounts the rate of positive-sense production (Table 1 reaction 6).

After newly synthesized positive-sense genomes are produced, they can either be packaged into new virion particles (Fig. 2A, step 7), diffuse into the cytoplasm and begin translation (Fig. 2A, step 8), or remain in the current complex to continue replicating. The probability of packaging a newly synthesized positive-sense genome (Fig. 2A, step 7) is a function of a parameter that describes the rate of packaging (*c*_*pack*_) and the amount of available capsule proteins (*CAP*). The probability of a newly synthesized positive strand dispersing in the cell (Fig. 2A, step 8) is a function of the probability that it stays within the replication complex (*c*_*stay*_) or that it is packaged (*p*_*pack*_).

### Fitting the model of intercelluar replication dynamics to population-level data

[18] fitted their stochastic model of PV infection to population-level quantitative RT-PCR data of positive-sense and negative sense strands gathered from replicate experiments at six time points and three different multiplicities of infection (MOI). The fitting was performed via Approximate Bayesian Computation (ABC). Here, we modify this approach in two crucial aspects. First, instead of fitting the model to RT-PCR data ([18]’s approach), we fit the model to fluorescence time courses generated during infection. Second, instead of fitting the model to population-level data, we fit it to data representing individual infections in single cells. Since we have thousands of individual infection time courses that display substantial variation among them due to the stochastic nature of both infection dynamics and cellular environments, we cannot fit an individual set of model parameters. Instead, we have to assume that each parameter is represented by a distribution characterized by a mean and a variance, and we fit these distributions to the population of infection time courses. In effect, we are converting [18]’s fixed-effects model into a random-effects model to capture the variation among individual infection time courses due to variation in internal state among different infected cells. Because this random-effects model presents a much increased parameter search space we employ a Genetic Algorithm (GA) for initial parameter optimization, followed by multiple rounds of the ABC procedure outlined by [18] to estimate posterior distributions of parameter values.

Fitting the stochastic model of intercellular replication dynamics to a population of infection time courses is implemented as follows: We first randomly generate a set of model parameters that represent the viral population that is to be simulated (Fig. 2B, step 1). This set of model parameters consists of a mean (*µ*_*i*_) and a variance (*σ*_*i*_) parameter for each of the model’s mechanistic variables. We then generate a population of individuals by randomly drawing sets of variables from the distributions specified by *µ*_*i*_ and *σ*_*i*_ (Fig. 2B, step 2) and simulating the infection dynamics for each individual (Fig. 2B, step 3). Each individual simulation produces an infection time course from which we extract the slope, maximum, midpoint, and time of lysis parameters as described by [17] (Fig. 2B, step 4). The estimates of these parameters are then pooled among individuals to create distributions that describe the population of infections (Fig. 2B, step 5). We then compare each of these four simulated parameter distributions to the experimentally observed distributions with a K-S test (Fig. 2B, step 6). The K-S tests produce a *D* statistic that represents how similar each of the four simulated parameter distributions are to the experimental observations. The smaller the value of *D* the more similar the simulated and experimental distributions are. We sum over the *D* values for each of four distribution comparisons to create a score for the population parameters (Fig. 2B, step 6). We then employ GA followed by multiple rounds of ABC using this scoring procedure to minimize the difference between the simulated and experimental parameter distributions to estimate the mechanistic parameters in our model of PV infection (for details, see Methods: Model Fitting).

### Estimates of inter-cellular replication dynamics

As the first application of our simulation model, we fit its output using a combination of GA and ABC to the distributions of the slope, maximum, midpoint, and lysis parameters extracted from single-cell infection time courses performed without any drug treatment, and we visually assess that our fitted distributions correspond to the experimental observations (Fig. S1A–D). We also examine Q–Q plots that compare the simulated and experimentally observed data to confirm the quality of fit (Fig. S2A–D). To assess whether our model behaves as a proxy for viral replication, we track when in time we first observe protein and RNA production in our model. We find the median amount of time until first protein production is close to 1 hour, while it takes approximately 4.5 hours for positive sense RNA production to be observed, and 5.5 hours until negative sense RNA production observed (Fig. 3). The temporal order and timing of these events is consistent with the known biology of PV replication [20].

**Figure 3:**
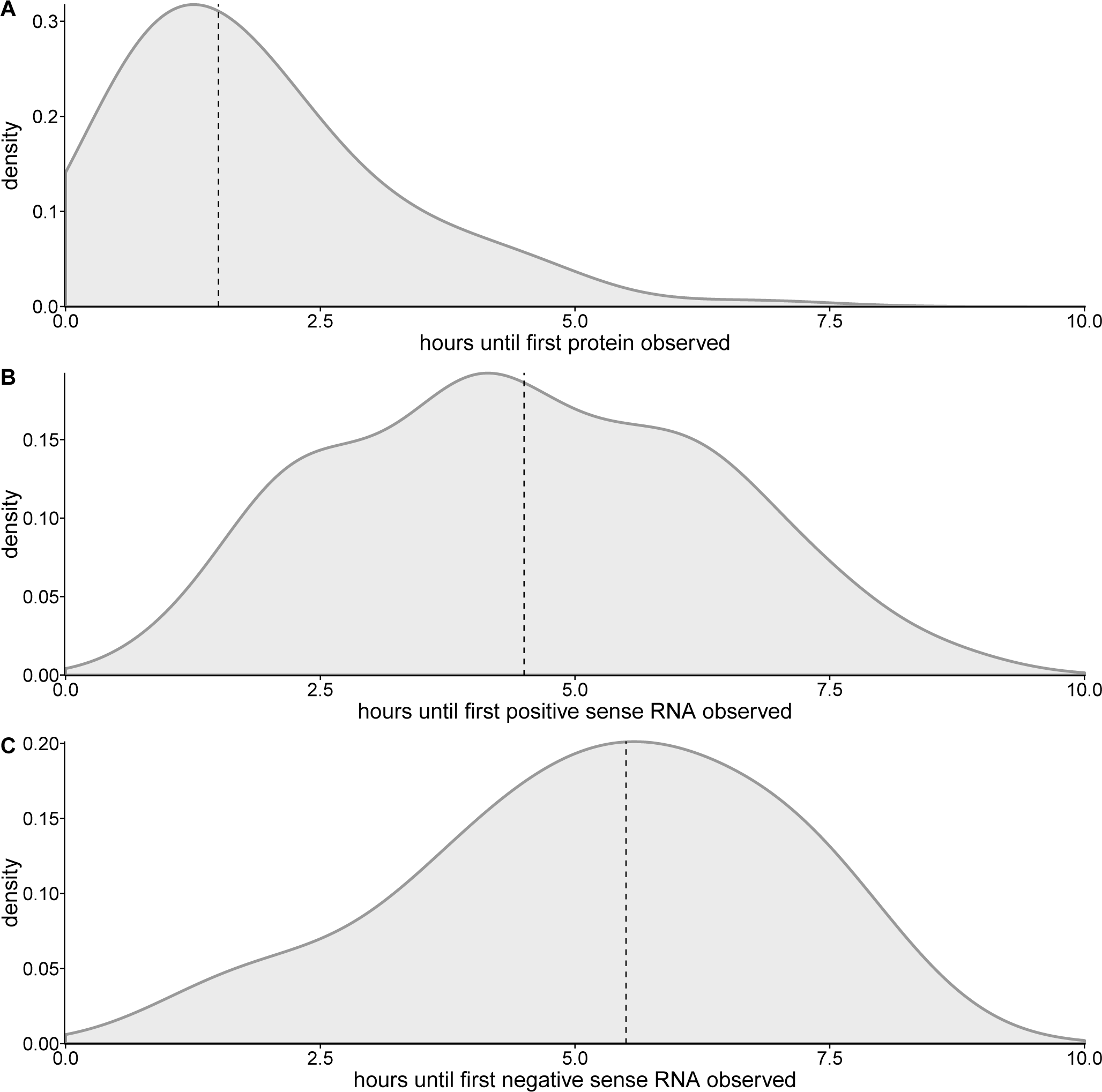
Time until first occurrence of events as estimated from our model of PV infection when the model is fit to the experimental data shown in Fig. 1C–F. (A) Hours until protein is first produced. (B) Hours until the first production of positive sense RNA. (C) Hours until the first production of negative sense RNA.

Next, we examine the posterior distributions of fitted parameters from our mechanistic model to discern which parameters are informative for describing experimentally observed population-level replication dynamics. We find that several of our posterior distributions appear to converge, while others do not (Fig. 4). Parameters whose posterior distributions converge include our estimates of the parameters that describe the processes of translation (*c*_*trans*_) (Fig. 4A), complex formation (*c*_*com*_) (Fig. 4B), circularization (*c*_*circ*_) (Fig. 4C), and positive strand replication (*c*_*rep*_+) (Fig. 4D). Estimates of the posterior distributions of these parameters were observed to converge by [18] as well. Our observation that these distributions appear to converge suggests that individuals within a viral population behave similarly in regards to translation, compartmentalization, circularization, and positive strand replication. Further, appropriately capturing these dynamics appears to be important for our model to describe experimentally generated population-level growth data.

**Figure 4:**
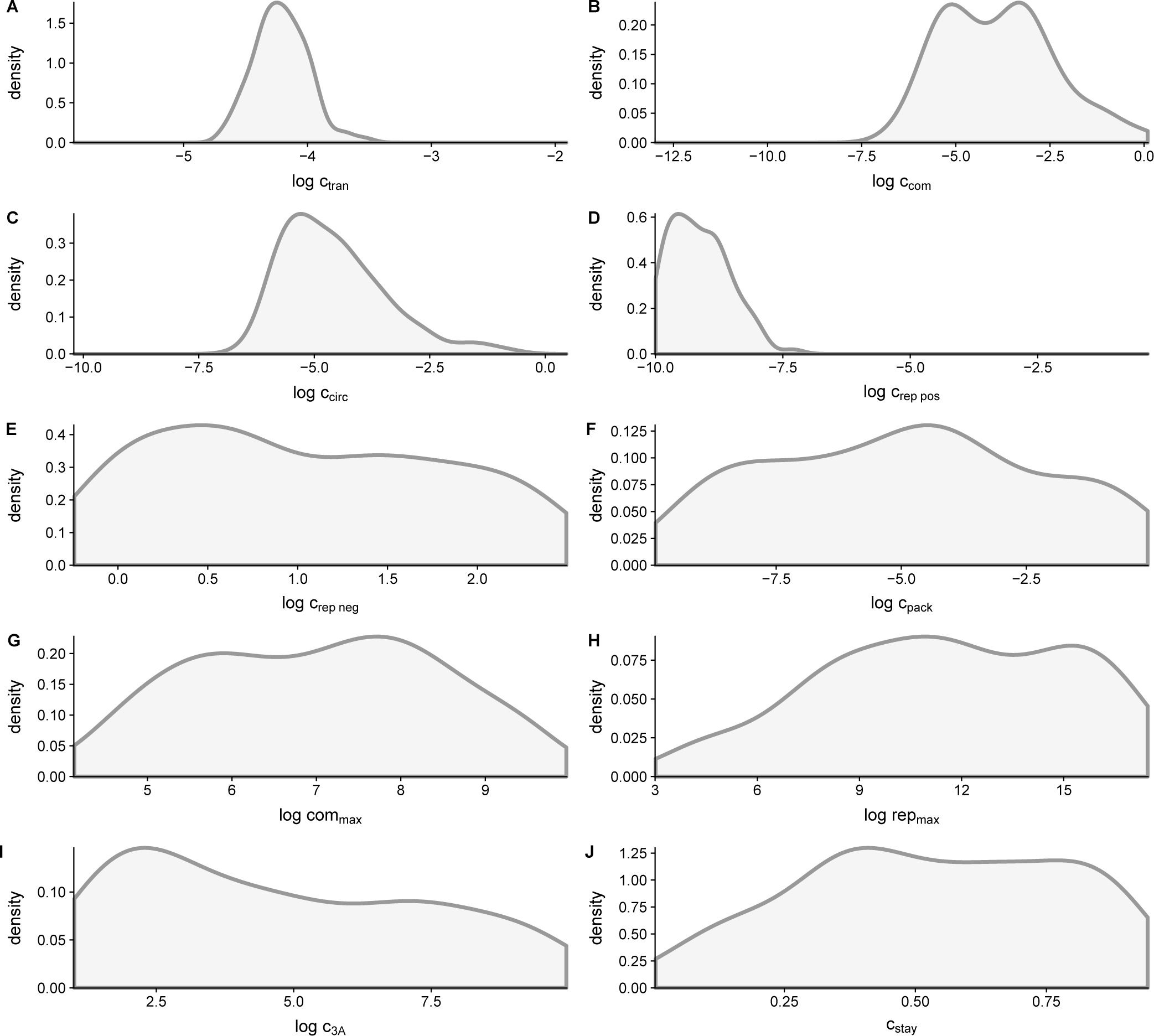
Estimates of mechanistic parameter posterior distributions in our model of PV infection from fitting the model to experimental data generated without drug treatment (Fig. 1C-F). Parameters are estimated by fitting the model described by the equations in Table 1 and illustrated in Fig. 2A. The parameters shown correspond to those labeled in each reaction. (A) Translation, which occurs in step 3 of the model. (B) Compartmentalization, a part of step 4. (C) Circularization, step 5. (D, E) Replication of positive and negative sense RNA, step 6. (F) Packaging, step 7. (G) The maximum number of compartments possible, considered in step 6. (H) The maximum number of replication cycles permitted by cellular resources, a limiting factor in step 4. (I) Consumption of the protein product 3*A*, step 4. (J) The probability for a newly synthesized genome to stay in the replication complex, step 8.

Posterior distributions for parameters associated with negative strand replication, packaging, and the maximum number of compartments (Fig. 4D–G) have large variance. This variation in posterior estimates suggests that PV may exhibit variability in how the respective processes are being carried out within individual cells, or that these aspects of the model are less essential for capturing dynamics observed in the experimental data. We observe skewed posterior distributions of our estimates of the maximum number of replication cycles (*rep*_*max*_) (Fig. 4H), consumption of the protein product 3*A* (Fig. 4I), and the probability of a newly synthesized positive strand to stay within the replication complex (Fig. 4J). These skewed distributions may suggest that while variation exists, an increased maximum number of replication cycles, a reduced amount of protein consumption, and a high probability of a new strand to stay in the replication complex occur more often.

### Informative parameters

To further examine which parameters in our model are informative for estimating experimentally observed growth data we perform principal component analysis on the posterior estimates. We find that the first principal component explains nearly 18% of the observed variance, and that the first six principal components explain a cumulative 73% of the observed variance (Fig. 5A). Examining the relative contribution of features in each of the principal components, we find that the parameters that describe complex formation and circularization have the largest amount of relative contribution in the first principal component (Fig. 5B). These two parameters are expected to be related since circularization occurs within complexes (Fig. 2B steps 4 and 5). Considering these two features in the first principal components are also parameters we observed to converge (Fig. 4) suggests that these parameters are an important aspect of our model for generating viral growth data.

**Figure 5:**
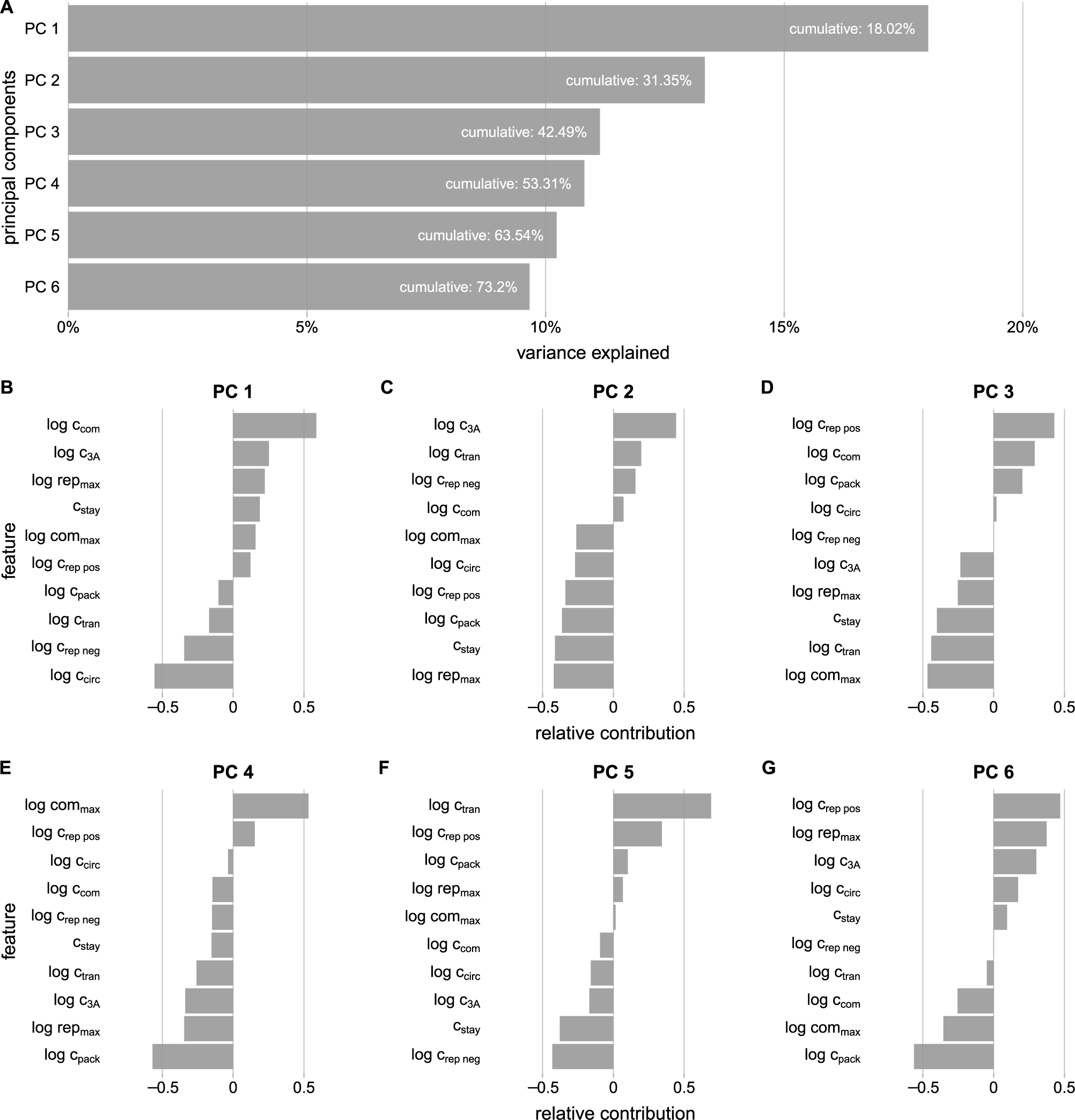
Principal component analysis of parameter distributions estimated from fitting our mechanistic model of PV infection to experimental data (Fig. 1C–F). (A) The amount of variance explained by the first 6 principal components. The white labeling provides the cumulative amount of variance explained by including all components up to and including the labeled one. (B–G) Relative contribution of features for each of the first six principal component axes.

The maximum number of replication events (*rep*_*max*_) and the consumption of the protein product 3*A* have the largest amount of relative contribution in the second principal component (Fig. 5C). In the model the maximum number of replication events represents an upper bound on replication, and it may be influential on the maximum amount of GFP produced in our simulations. The protein product 3*A* is needed for complex formation, suggesting that accounting for the use of resources produced during translation may also be an important aspect of modeling viral growth data.

Additionally, to investigate how parameters interact within our model we examine the correlations between parameters (Fig. S3). We find that the strongest relationship exists between parameters associated with compartmentalization and circularization. Interestingly, these parameters are found to have the largest amount of relative contributions in the first principal component (Fig. 5B). Other parameter estimates are less correlated, though there appears to be a relationship between parameters associated with the maximum number of replication cycles and the probability that a new strand will stay in its replication complex, as well as translation and complex formation.

### The effect of drug treatments on parameter estimates

To examine if the mechanisms found to be informative about viral growth dynamics in our initial data set—without drug treatment—also describe the variability observed in experiments with drug treatment, we expand our analysis to three data sets generated using different classes of drugs (Fig. S1E–P, Fig. S2E–P). We find that the estimates of several of our mechanistic parameters differ when fitting our model to these additional three data sets. Treatment with the drug rupintrivir, a 3C protease inhibitor, affects our estimates of parameters associated with translation, compartmentalization, circularization, and replication of positive strands (Fig. 6A–D). Rupintrivir increases the range of parameter estimates we find to be informative when fitting data generated without drug treatment. However, when fitting the model to experimental data generated under treatment with rupintrivir we observe poor convergence (Fig. S1E–H, Fig. S2E–H), though it is notable that only the posterior distributions of the four parameters that clearly converge in the no-drug case differ.

**Figure 6:**
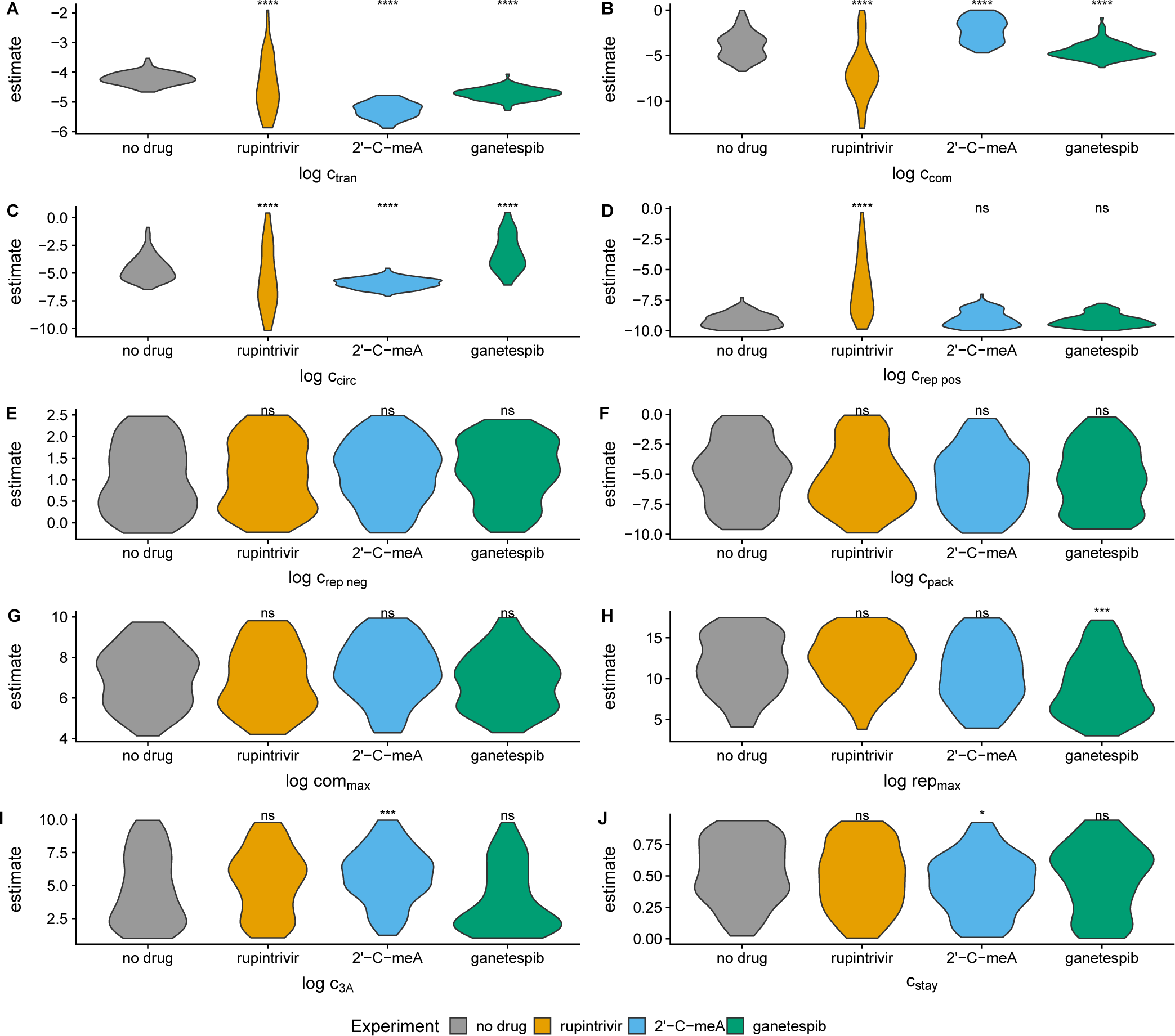
Comparison of posterior parameter distributions between the no drug treatment and drug treatments. Asterisks indicate distributions that differ significantly from the no drug treatment. Non significant (ns) corresponds to *p* > 0.05, a single asterisk corresponds to *p* ≤ 0.05, two asterisks correspond to *p* ≤ 0.01, three correspond to *p* ≤ 0.001, and four correspond to *p* ≤ 0.0001 from a K-S test. Parameters are estimated by fitting the model described by the equations in Table 1 and illustrated in Fig. 2A. Parameters correspond to those labeled in each reaction. (A) Translation, which occurs in step 3 of the model. (B) Compartmentalization, a part of step 4. (C) Circularization, step 5. (D, E) Replication of positive and negative sense RNA, step 6. (F) Packaging, step 7. (G) The maximum number of compartments possible, considered in step 6. (H) The maximum number of replication cycles permitted by cellular resources, a limiting factor in step 4. (I) Consumption of the protein product 3*A*, step 4. (J) The probability for a newly synthesized genome to stay in the replication complex, step 8.

Fitting our model to experimental observations under treatment with 2’-C-meA, a polymerase inhibitor, we find that our estimates of parameters associated with translation, compartmentalization, circularization, consumption of protein products, and the probability of a new strand staying in its replication complex differ (Fig. 6A–C, I, J) (see also Fig. S1I–L, Fig. S2I–L). 2’-C-meA decreases the translation (Fig. 6A) and circularization (Fig. 6C) parameters, while increasing the parameters associated with compartmentalization (Fig. 6B) and the consumption of 3*A* (Fig. 6I). Notably, our estimate of the probability of a new strand to stay in the replication complex is decreased under this drug treatment (Fig. 6J). 2’-C-meA is the only drug treatment where we find a substantial change in the probability of a strand to stay in its replication complex, suggesting that this treatment impacts the viral replication process differently than the other two treatments.

Fitting our model to experimental observations under treatment with ganetespib, an Hsp90 inhibitor, impacts our estimates of translation (Fig. 6A), compartmentalization (Fig. 6B), cicularization (Fig. 6C), maximum number of replication cycles (Fig. 6H), and the probability of a new strand staying in its replication complex (Fig. 6A–C, H, J) (see also Fig. S1M–P, Fig. S2M–P). Treatment with ganetespib appears to similarly impact our estimates of some aspect of the protein production and utilization process as the other drug treatments. The most notable change to parameter estimates under treatment with ganetepib is the decrease in the maximum number of replication cycles (Fig. 6H). This treatment is the only out of the three to do so. In aggregate, we find that each of the drug treatments affects different parameter estimates. This finding suggests that treatment with each of the drugs impacts the viral replication process differently.

The drug treatment data also result in estimates of parameter variances that differ from the case of no-drug treatment in several cases (Fig. S4). Each of the drug treatments appears to impact variation in translation (Fig. S4A) and in the maximum number of replication cycles (Fig. S4H). Rupintrivir also affects variation in the probability that a new strand will stay in its replication compartment (Fig. S4J), while ganetespib affects variation in the maximum number of compartments (Fig. S4G).

To examine which parameters are informative for estimating experimentally observed growth data under drug treatments we again perform principal component analysis. We find that the first 6 principal components of each drug treatment explain around 75% of the variance (Fig. S5A, D, G), similar to that found in the no drug case. Examining the features of the first and second principal components under treatment with rupintrivir, we find that parameters associated with compartmentalization, strand replication, and packaging have the largest relative contributions (Fig. S5B, C). The first and second principal components under treatment with 2’-C-meA show that parameters associated with consumption of the protein 3*A*, circularization, the maximum number of compartments and replication cycles have the largest relative contributions (Fig. S5E, F). When fitting our model to experimental data treated with ganetespib we again find parameters associated with circularization, compartmentalization, strand replication, and packaging also have the largest relative contributions (Fig. S5H, I).

Notably, many of the parameters found to have a larger amount of relative contribution in the first two principal components of each of the drug treatments are found to have larger relative contributions in the no drug case. Considering that parameters associated with compartmentalization and circularization have a larger amount of relative contribution in the first two principal components in many of our experiments suggests that these parameters are important for our model to recapitulate experimental observations.

Additionally, drug treatments influence the correlations between parameters. Treatment with rupintrivir results in a negative correlation between compartmentalization and the replication of positive strands (Fig. S6). Treatment with the other two drugs 2’-C-meA (Fig. S7) and ganetepib (Fig. S8) results in a correlation between compartmentalization and circularization, similar to that observed in the no drug case (Fig. S3).

### Estimates of the number of generations between infecting and packaged virions

[18] used their model to estimate the mean number of generations between infecting and packaged virions. We repeat this analysis on our four data sets to assess if our parameter estimates generate similar generation estimates (Fig. S9). Examining the distributions of mean number of generations between infecting and packaged virions, we find that these distributions are slightly right skewed and two of the drug treatments generate distributions that are significantly different from the no drug case (Fig. S9B, C). The long tails of these distribution suggest that a small fraction of packaged virions undergo an increased number of generations. The median estimate of the mean number of generations in the treatment without the drug is 4.75, while treatment with rupintrivir results in 5.77 generations, treatment with 2’-C-meA results in 4.67 generations, and treatment with ganetespib results in 4 generations (Fig. S9). We find that rupintrivir (Fig. S9B) and 2’-C-meA (Fig. S9C) both increase the median of the distribution of generations for a packaged virion compared to the no drug case, while treatment with ganetespib decreases the median number of generations (Fig. S9D).

The number of replication cycles in PV has previously been estimated to be 2.1–4.6 [21], and these estimates were based on measurements of the initial and final virus titers combined with burst size measurements taken from previous studies [2, 22–24]. [18] estimated a slightly higher number of approximately 5 replication cycles. Our estimate of between 4–5.77 replication cycles seems to correspond well to the estimates from these other studies, suggesting that, similarly to our findings regarding the timing of events (Fig. 3), our model is behaving as a reasonable proxy of biological reality.

## Discussion

We have developed a model-fitting framework to estimate parameters of viral replication by fitting a stochastic model of PV replication in a single cell to a population of infection time-courses measured in individual infected cells. Using this framework, we have identified the features of the model that are critical for capturing viral growth dynamics. We find that accounting for translation and for early stage processes such as complex formation and circularization are the most essential aspects of the model. Estimates of the parameters representing these processes appear to converge and are found to be important features in PCA analysis. We have also confirmed that the model captures several biologically relevant aspects of viral replication not obviously visible in the input data, such as the timing of intercellular events and the number of generations between infecting and packaged virons.

Using three experimental data sets generated under various drug treatments, we have observed substantially different estimates of critical parameters compared to the no-drug case. The differences in the posterior distributions under drug treatment may be due to the fact that our model does not include a mechanistic description of these drug effects. Hence, our estimates of parameters from these experiments may not necessarily reflect how specifically drug treatments impact aspects of viral replication, but rather they may highlight which parameters in our model are capable of capturing the variation in growth dynamics under various experimental conditions. Additionally, we have found that two of the drug treatments, rupintrivir and 2’-C-meA, lead to an increased median number of generations between infecting and packaged virions. This observation is most likely due to the fact that drug treatments lead to a longer time until cell lysis, increasing the opportunity for more generations to occur.

The observation that some drug treatments increase the length of time until cell lysis generates the interesting hypothesis that some drug treatments may increase the number of replication cycles inside the cell. The increased number of replication cycles would indicate a greater effective mutation rate under some drug treatment. Due to the relationship between the number of replication cycles, mutation accumulation, and drug resistance, examining the number of replication cycles is critical for developing effective and safe antiviral treatment regimens, and therefore many studies attempt to estimate it [14, 25– 27]. Drug treatments have been shown to affect genetic diversity in various other systems. Treatment with 3-azido-3-deoxythymidine (AZT) has been found to influence the *in vivo* mutation rate of HIV-1 and can increase the rate of mutation by a factor of 7 in a single round of replication [28]. Treatment with favipiravir has been shown to result in an effective increase in the viral mutation rate of influenza A virus [29]. The broad-spectrum antiviral drug ribavirin is an RNA virus mutagen that can cause a 9.7-fold increase in mutagenesis in poliovirus resulting in error catastrophe [30]. Drug treatments have also been shown to increase viral replication. Corticosteroids increase viral replication in the adenovirus type 5/New Zealand rabbit ocular model [31]. Valproic acid A has been shown to increase viral replication in HIV [32]. Treatment with rapamycin or everolimus facilitates hepatitis E virus replication [33]. Extending the model used here to account for the mechanisms of a drug’s action to examine how drug treatments impact the PV replication process could be a useful step for studying the relationship between drug treatments and viral evolution.

Our observations of which aspects of the viral replication process are important for capturing experimentally observed viral dynamics are based on a specific model, and our findings are only sound if the model is indeed capturing biological reality. Considering that our prediction of the timing of events and approximately 4–5.77 replication cycles align with the previous literature, we believe that the model correctly captures at least some aspects of the viral replication process. To further improve the model, a concise picture of how the replication process occurs is needed. Results from this work also highlight the need for more quantitative experimental approaches to enhance model fitting. For example, if the formation of replication complexes could be microscopically tracked, including this data in the fitting process could be beneficial. Further work is necessary to expand the model to encompass greater levels of biological reality. To explore intracelluar dynamics in greater detail this model could be extended to include more detailed description of each of the reactions in the model along with parameters that account for intracellular selection, recombination, and mechanisms of the effects of drug treatments. However, extending the model does have some drawbacks. Accounting for further levels of biological reality would require an increase in the number of parameters in the model, which could lead to overfitting. It would also likely increase the amount of computational time necessary for the model to converge.

Fitting stochastic models to population-level time-course data is a relatively new field of research. One issue that consistently arises in this field is that fitting by comparison of experimental and simulation distributions has the distinct drawback of high computational execution time [34–37]. We have encountered this issue in our study as well. The ABC runs can take several weeks to converge for a single set of experimental data, even when the fitting code takes advantage of parallel model evaluations running on over 100 CPUs at the same time. In other words, convergence of a single ABC run takes on the order of 20,000 to 50,000 CPU hours. This high computational cost arises because we need to execute hundreds of independent simulation runs to evaluate any one given set of parameter values. Because the model is already so computationally costly, expanding it to incorporate further levels of biological reality would undoubtedly increase the run time even further. Thus, irrespective of the specific analysis we have carried out here, we see a need to develop more efficient algorithms for fitting stochastic models to population level data.

In conclusion, the results presented here represent a stepping stone in the pursuit of using mechanistic modeling approaches to tease apart details of high-throughput single-cell viral growth dynamics that may be unobservable with experimental approaches alone. Additionally, this work introduces a framework for fitting models designed to describe single time-course trajectories to population level dynamics. Models like these can inform researchers about which aspects of the viral replication process are more variable or stable than others. Identifying which aspects of intercelluar dynamics lead to the variation observed in viral growth data could lead to insight relevant to drug development efforts. Though our findings are limited by the biological reality that our model is able to capture, they suggest that factors that occur early in the viral replication process are a key aspect of describing viral growth data.

## Methods

### Processing Input Data

Using previously published data [12] from single-cell PV growth experiments generated under four conditions, no drug, 2’-C-meA treatment, rupintrivir treatment, and ganetespib treatment, we compile several metrics prior to model fitting. First, for each of the experimental conditions, we only retain infection time courses that can be classified as either sigmoidal or double-sigmoidal according to the sicegar R package [17]. Using the estimates of slope, midpoint, maximum, and lysis time produced by sicegar, we generate the parameter distributions that describe each of the experimental conditions. In cases where the cell fails to lyse within 24 hours, we record the lysis time as 24 hours.

### Model Fitting

Our model fitting and analysis procedure is implemented in the R package spire (Single-cell Population Infection Replication Estimation) (see Code and Data Availability below), which builds on PV simulation code originally developed by [18].

Our model of PV replication consists of 25 parameters. These parameters include 10 mechanistic parameters and their variances that describe infection dynamics within a cell, a florescence scaling term, the shape and rate parameters of a Gamma distribution of the time until lysis, and a lag time and its variance until florescence. We then sample from each of these distributions of 25 parameters to create a parameter vector, which we refer to as the instantiating vector. For each instantiating vector we generate 100 variations of the vector, by sampling from distributions determined by the instantiating vector’s parameter values. For each of the 100 variant parameter sets, we then simulate a PV infection time course using the model introduced by [18]. Each time course data produced by the model is then fit using the R package sicegar [17], which estimates the time of lysis, the maximum intensity, the midpoint, and the slope. We aggregate the estimates from each of 100 time courses to produce distributions of time of lysis, maximum value, midpoint, and slope. Finally, we compare our estimates of these distributions to the observed experimental distributions with a scoring function comprised of summing over the *D* values of Kolmogorov-Smirnov tests. The smaller the sum of the score, the better the instantiating vector describes the data.

To speed up convergence of our fitting procedure, several checks are run on a proposed parameter set before the PV simulation is run. If the bounds between the estimated lag and lysis time are outside of those experimentally observed, the parameter set is considered invalid. To prevent running PV simulations with excessively long run times, a timeout function is also implemented. If a single instance of the PV simulation takes longer then 5 seconds to run, the parameter set that spawned that instance of the simulation is considered invalid. When a time course is returned from the PV simulation, if sicegar is unable to fit the data then the parameter set is considered invalid. Additionally, if the estimates returned from sicegar of the slope, maximum, midpoint, and lysis are above or below the maximum and minimum experimentally derived values by more than 10% the parameter set is considered invalid.

To estimate the model parameters we first use a genetic algorithm (GA) [38] to quickly sample the range of valid parameter values. This GA is run for 100 generations, and the population of instantiating solutions of the final generation is then used to initialize a subsequent approximate Bayesian computation (ABC) fit. We employ the ABC parameter inference method introduced by [18] and run the ABC for numerous rounds. The completion of a round is determined by the algorithm accepting 100 instantiating parameter sets. At each round of the ABC we calculate the mean of the score function for the 100 accepted parameter sets and use the mean as an acceptance criterion during the next round of the ABC. This process of accepting 100 parameter sets and adjusting the acceptance criteria is repeated until the acceptance criterion drops below a value of 0.55, which takes approximately 15 rounds of ABC.

To record the timing of events in the PV simulation, we record the timing of each individual reaction in a simulation using the converged parameter set. From these timing records, we can identify when protein and RNA production takes place. Similarly, we can estimate the mean number of generations between infection and producing packaged virions.

### Data Analysis

Once the ABC acceptance criterion has dropped below the cut off we visually compare our estimated distributions of slope, maximum, midpoint, and lysis time to the experimental results (Fig. S1). To assess the correspondence of our estimated distributions to the experimental results we also examine quantile–quantile plots comparing the observed and estimated distributions (Fig. S2). From visual inspection and from examining the quantile– quantile plots, it appears that our estimated distributions correspond reasonably well to the experimental data, with the exception of fitting to data treated with rupintrivir which failed to converge after several weeks of running the fitting algorithm.

To examine how parameters within the model relate to one another, we calculate their correlation coefficients and cluster these correlations using hierarchical clustering. This clustering allows us to visualize related coefficients closer to one another. To examine which parameters are informative for estimating experimentally observed growth data we perform principal component analysis using the R package DataExplorer [39]. We compare the parameter estimates from the experiments under the no drug treatment to those estimated using drug treatments with a K-S test and we use a Bonferroni correction for multiple testing.

## Code and Data Availability

The software, results, and analysis tools, including the R package spire, are available at https://github.com/a-teufel/modeling single cell virology/. The repository has also been archived on Zenodo at https://doi.org/10.5281/zenodo.3695913.

## Acknowledgements

This work was supported by the National Institutes of Health grant AI120560 to C.E.C. and C.O.W.

---

**Figure S1:**
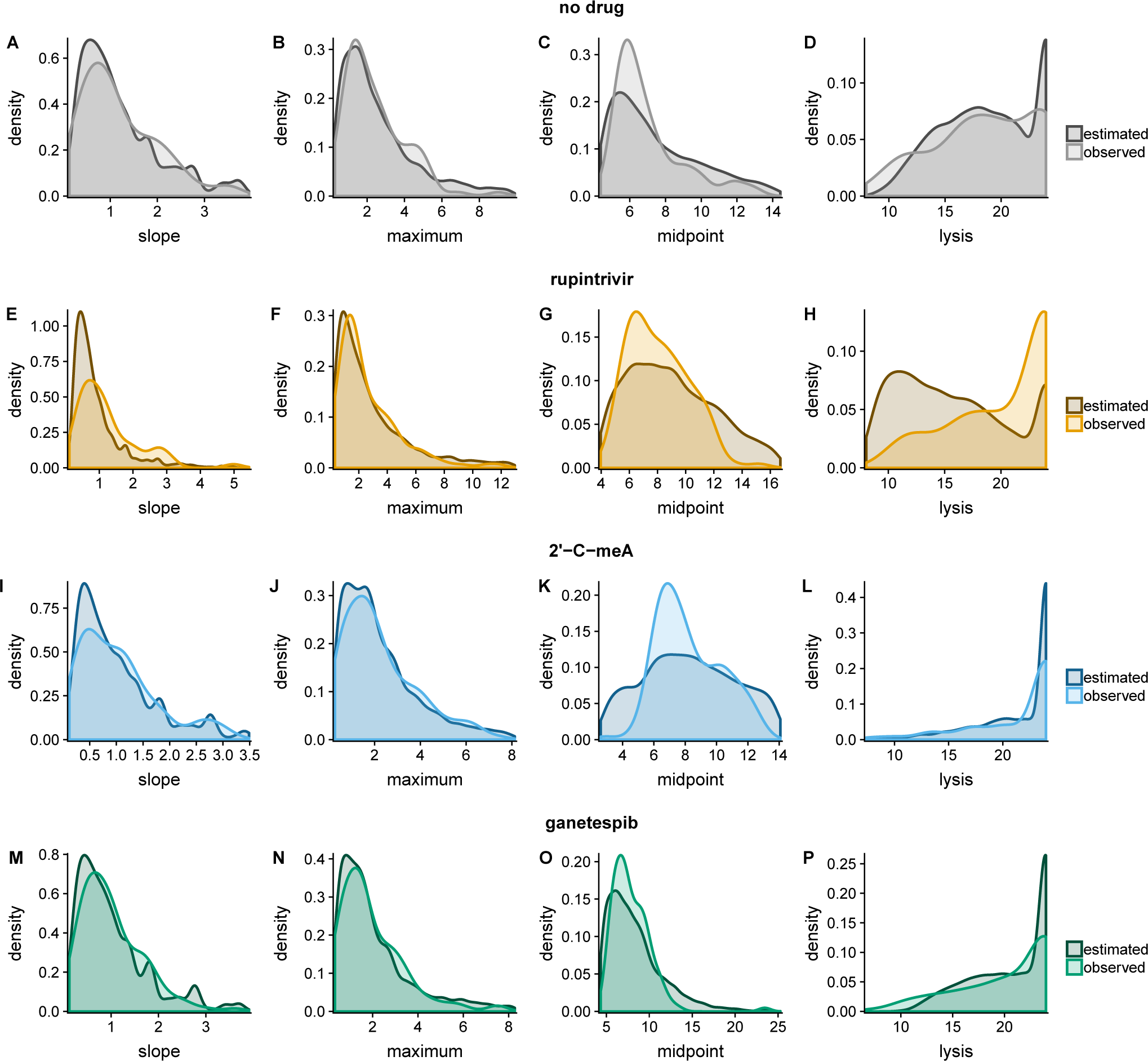
Visual comparison of estimated and experimentally observed distributions. (A–D) Results for experiments performed in the absence of drug treatment. (E-H) Results for experiments performed under treatment with rupintrivir. (I–L) Results for experiments performed under treatment with 2’-C-meA. (M-P) Results for experiments performed under treatment with ganetespib.

**Figure S2:**
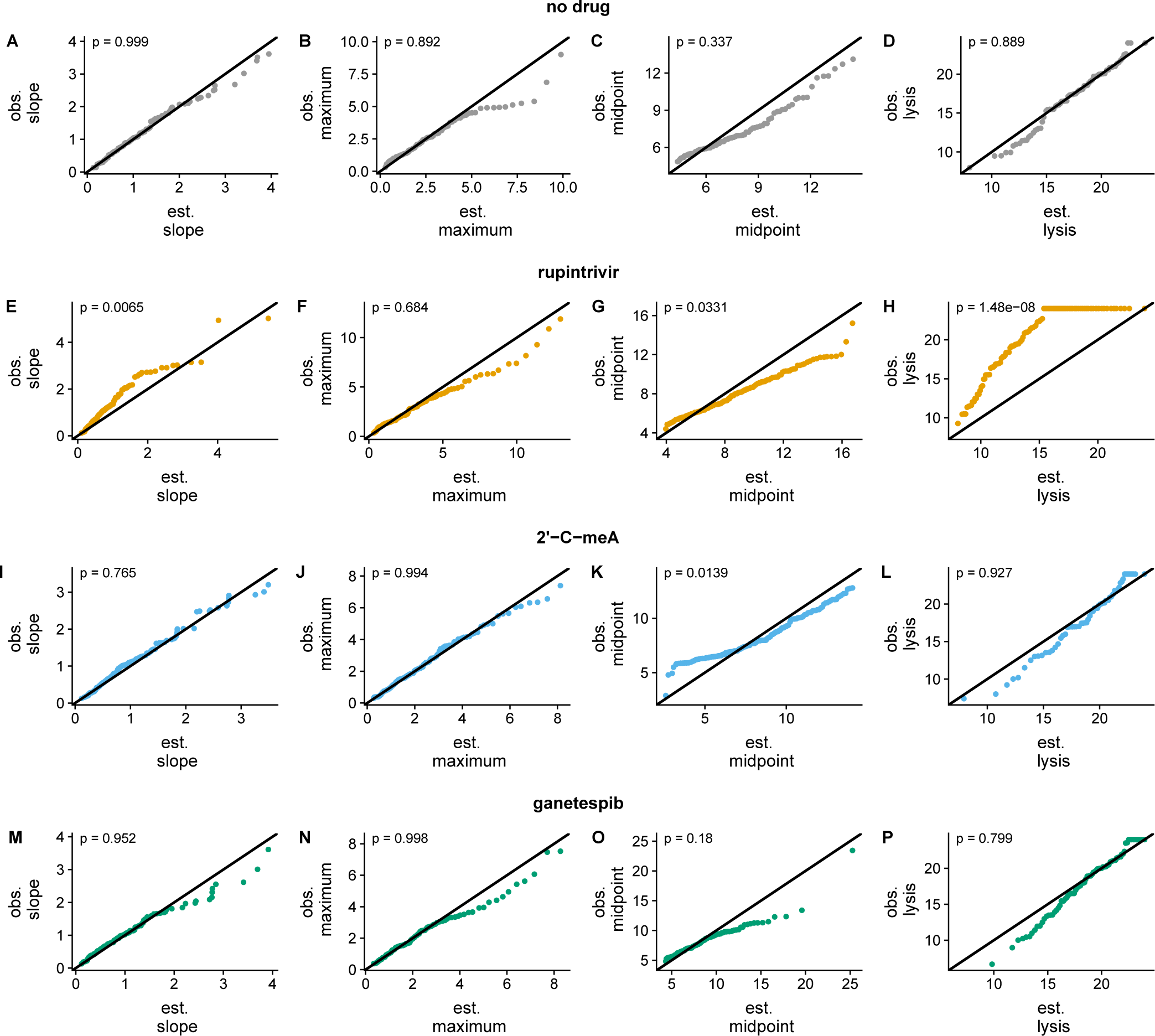
Quantile–quantile plots comparing estimated and observed distributions. *p*-values denote the results from K-S tests. (A–D) Results for experiments performed in the absence of drug treatment. (E-H) Results for experiments performed under treatment with rupintrivir. (I–L) Results for experiments performed under treatment with 2’-C-meA. (M-P) Results for experiments performed under treatment with ganetespib.

**Figure S3:**
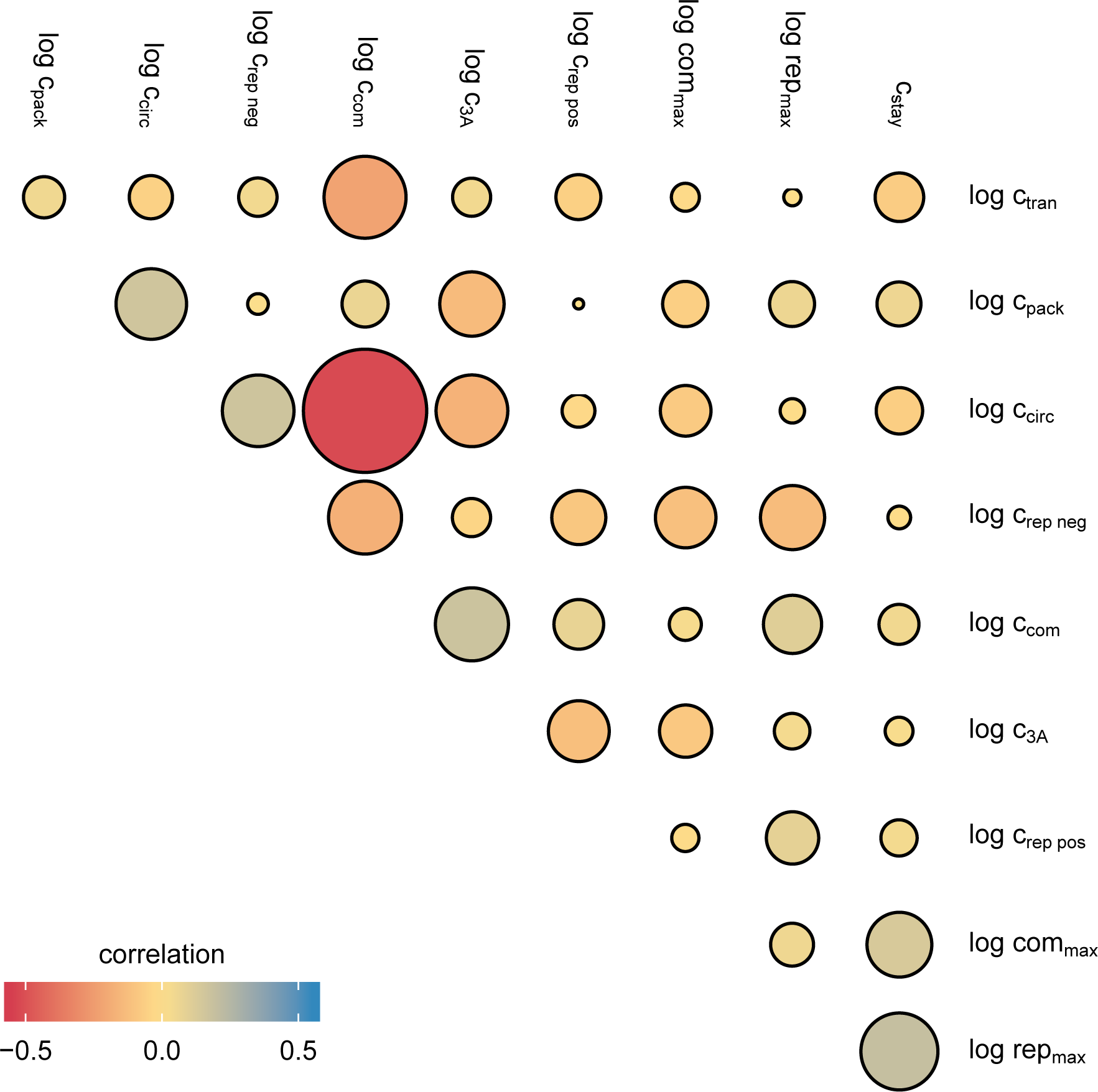
Correlations among parameter estimates from our mechanistic model of PV infection when the model is fit to the experimental data given in Fig. 1C-F.

**Figure S4:**
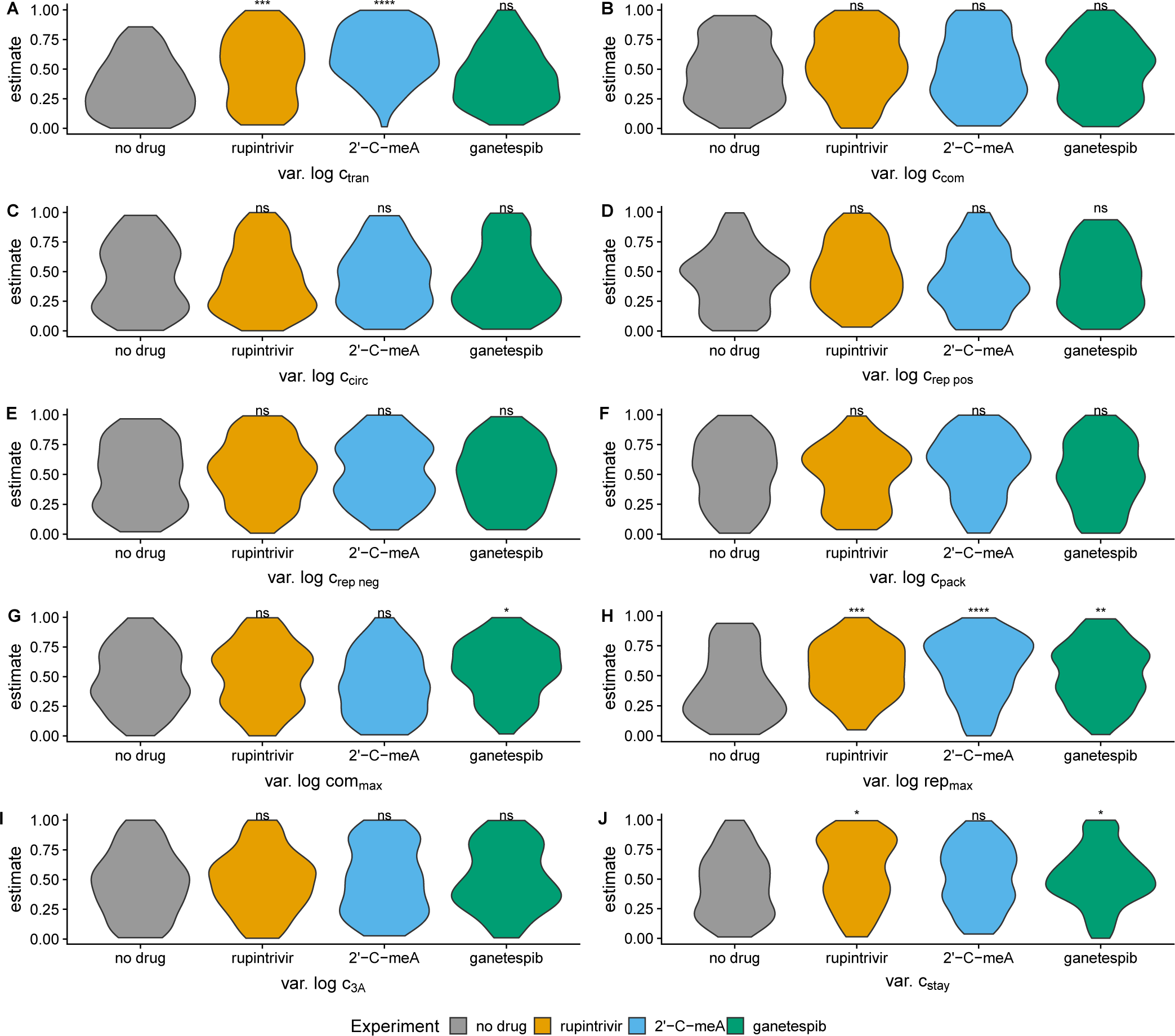
Comparison of posterior distributions for variance parameters between the no drug treatment and drug treatments. Asterisks indicate distributions that differ significantly from the no drug treatment. Non significant (ns) corresponds to *p* > 0.05, a single asterisk corresponds to *p* ≤ 0.05, two asterisks correspond to *p* ≤ 0.01, three correspond to *p* ≤ 0.001, and four correspond to *p* ≤ 0.0001 from a K-S test. Parameters are estimated by fitting the model described by the equations in Table 1 and illustrated in Fig. 2A. Parameters correspond to those labeled in each reaction. (A) Translation, which occurs in step 3 of the model. (B) Compartmentalization, a part of step 4. (C) Circularization, step 5. (D, E) Replication of positive and negative sense RNA, step 6. (F) Packaging, step 7. (G) The maximum number of compartments possible, considered in step 6. (H) The maximum number of replication cycles permitted by cellular resources, a limiting factor in step 4. (I) Consumption of the protein product 3*A*, step 4. (J) The probability for a newly synthesized genome to stay in the replication complex, step 8.

**Figure S5:**
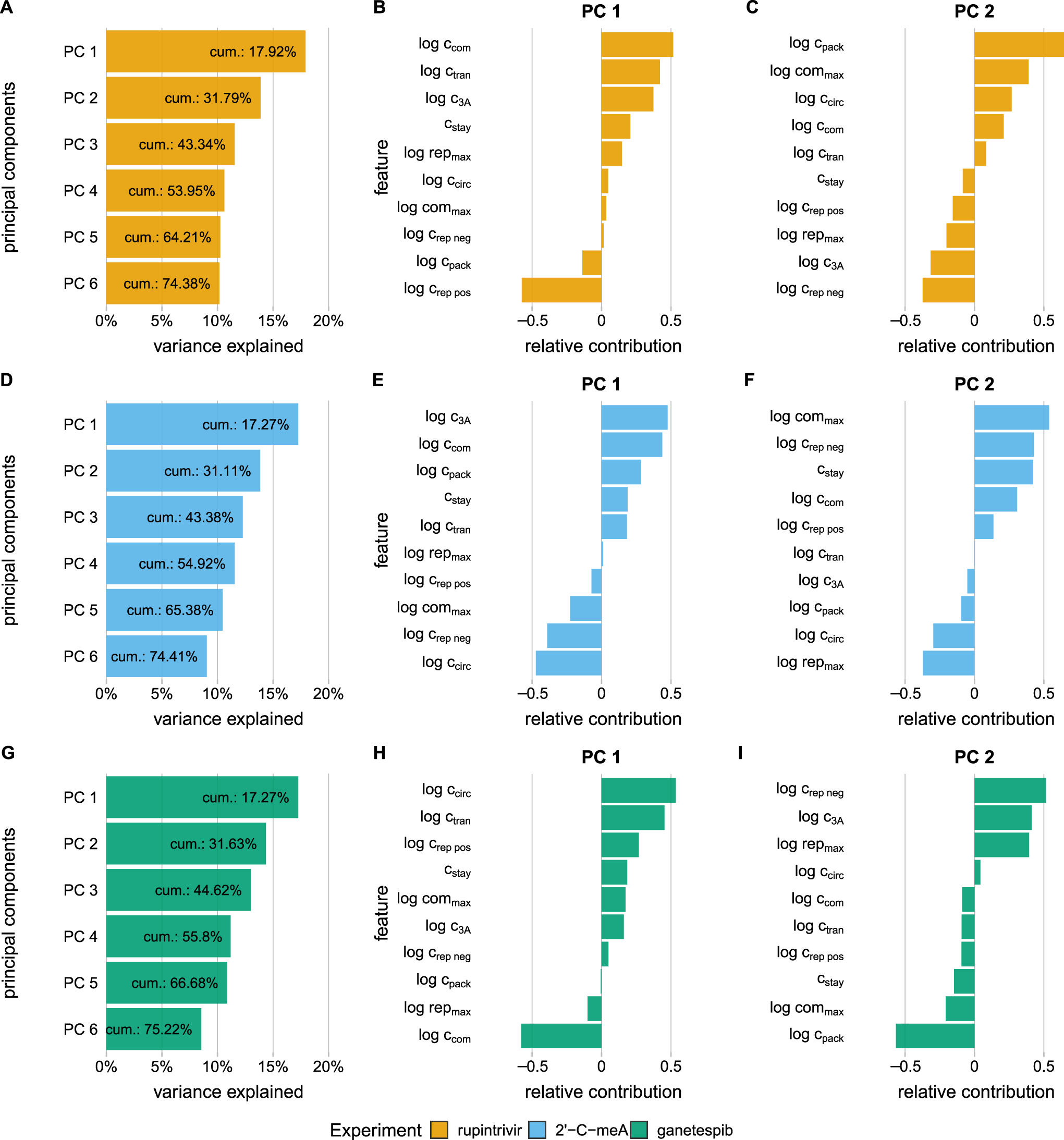
Principal component analysis of posterior parameter distributions estimated from fitting our mechanistic model of PV infection to experimental data generated under different drug treatments. (A) The amount of variance explained by the first 6 principal components when fitting to data generated under treatment with rupintrivir. Shown is the cumulative amount of variance explained by including each of the following components. (B, C) Relative contribution of features for the first and second principal component axis. (D) The amount of variance explained by the first 6 principal components when fitting to data generated under treatment with 2’-C-meA. Shown is the cumulative amount of variance explained by including each of the following components. (E, F) Relative contribution of features for the first and second principal component axis. (G) The amount of variance explained by the first 6 principal components when fitting to data generated under treatment with ganetespib. Shown is the cumulative amount of variance explained by including each of the following components. (H, I) Relative contribution of features for the first and second principal component axis.

**Figure S6:**
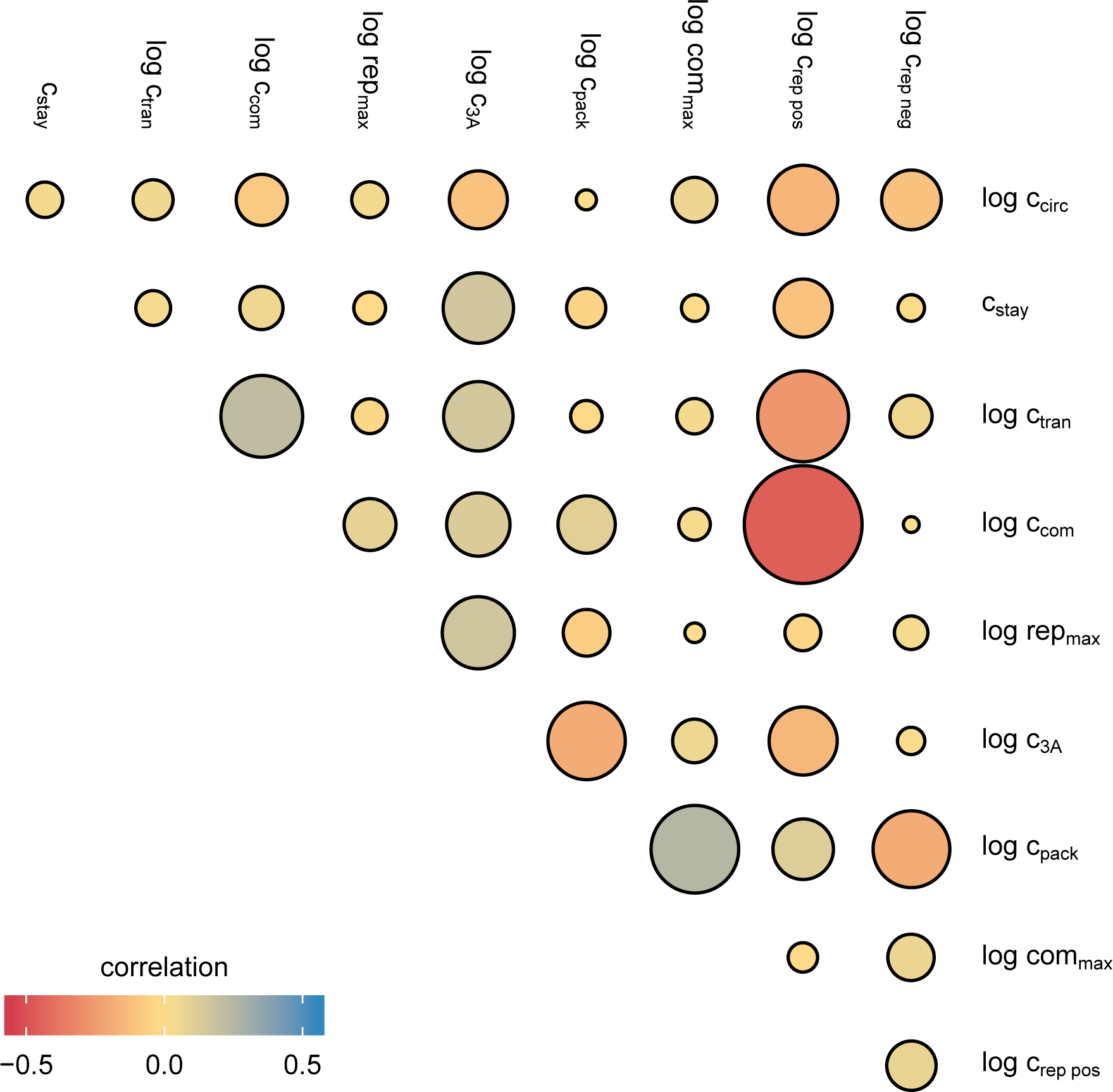
Correlation of parameter estimates when fitting data generated under rupintrivir treatment.

**Figure S7:**
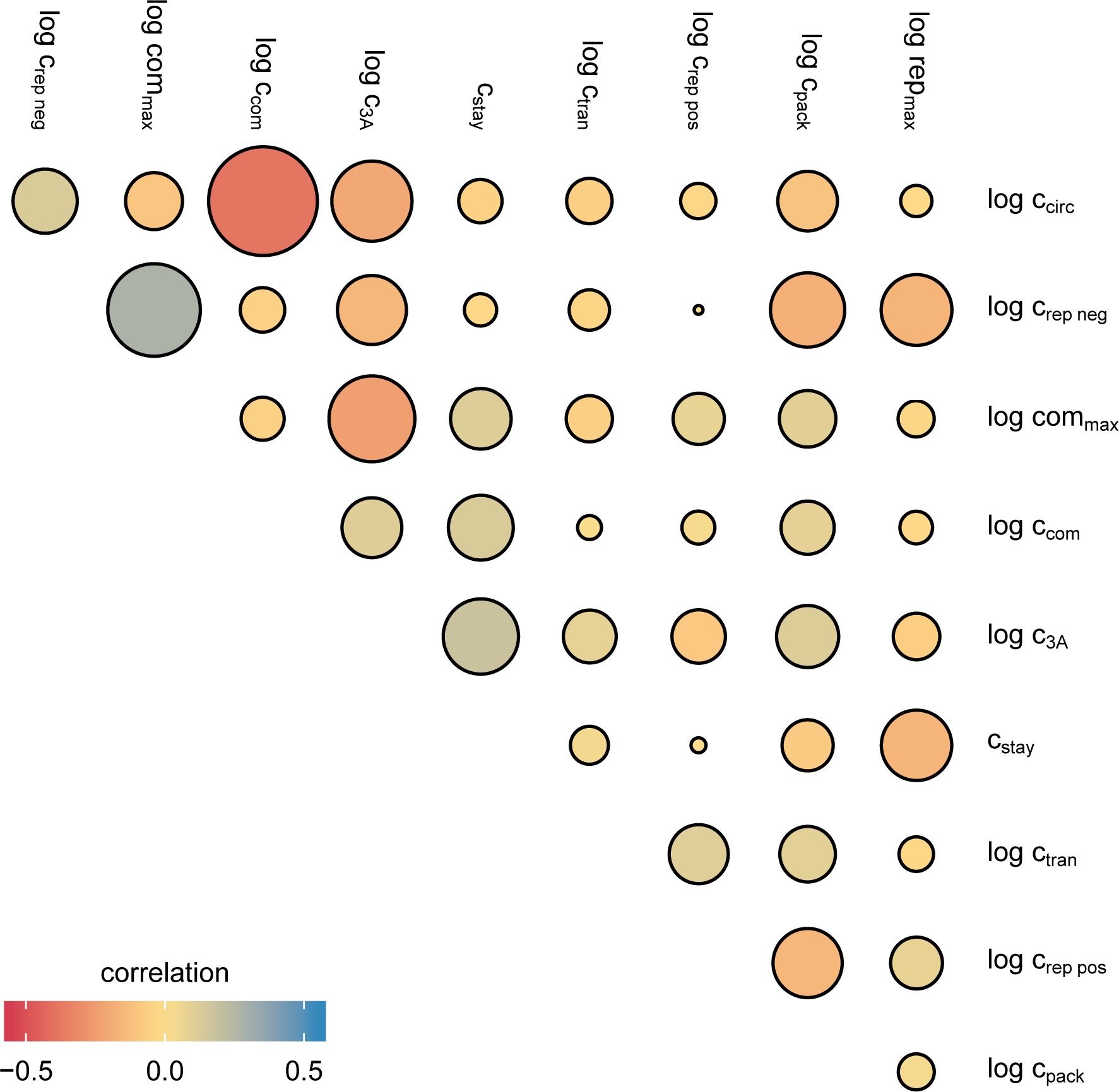
Correlation of parameter estimates when fitting data generated under 2’-C-meA treatment.

**Figure S8:**
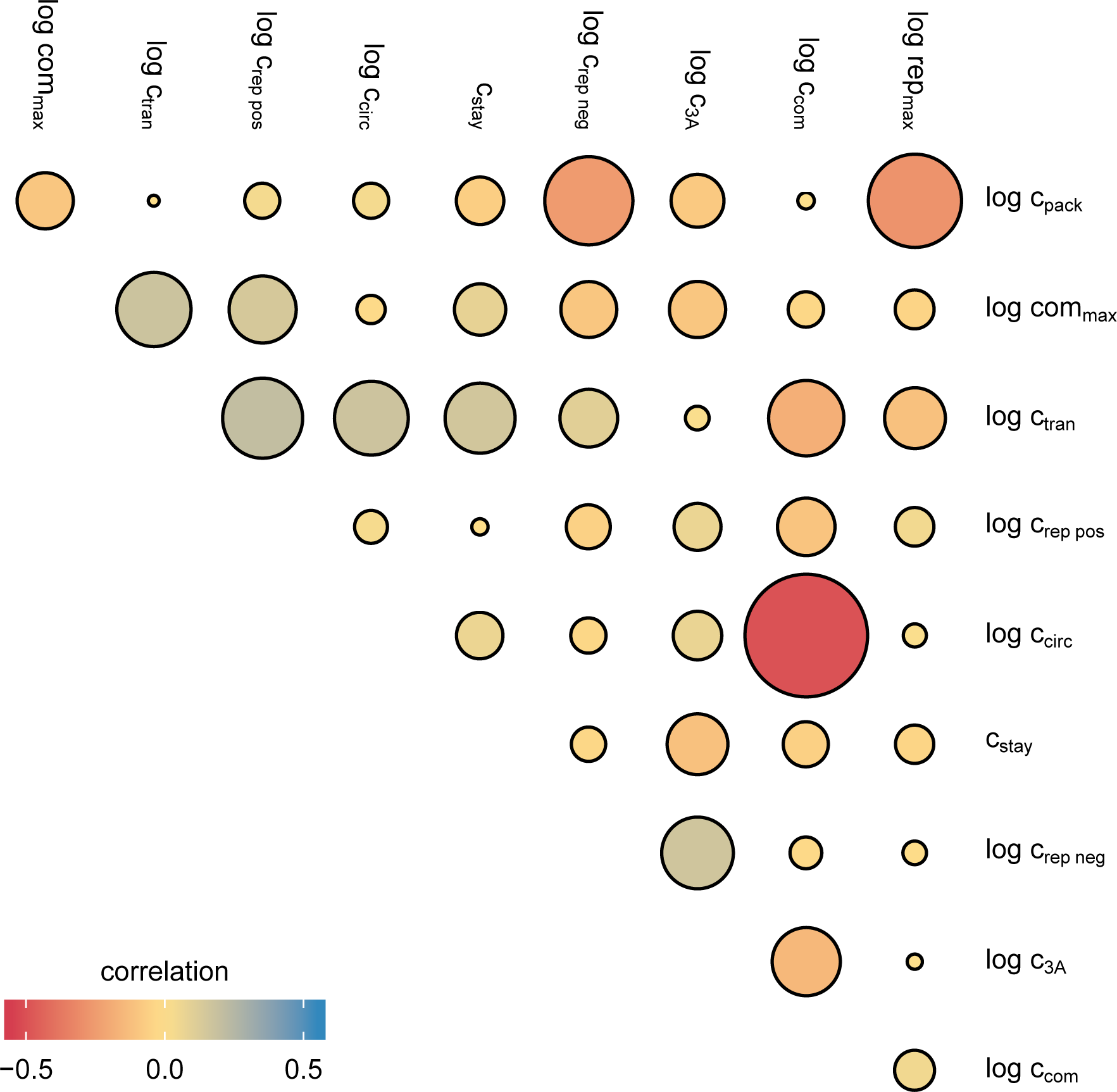
Correlation of parameter estimates when fitting data generated under ganetspib treatment.

**Figure S9:**
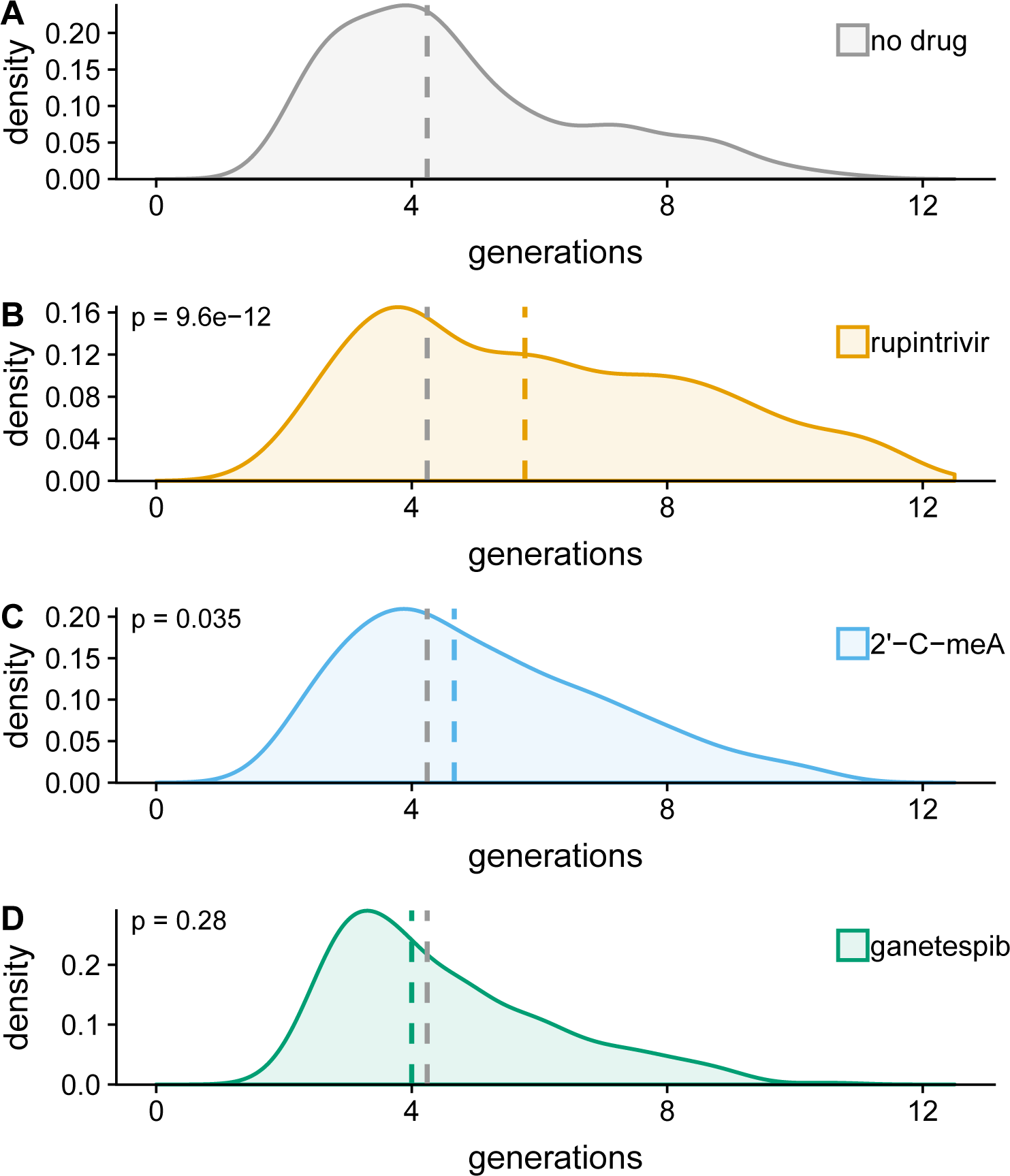
Comparison of mean number of generations under each of the experiments. Dashed lines denote the medians of the distributions. Gray dashed lines denote the median for the case of no drug treatment and are included in the drug treatment panels to allow for a comparison of the medians. *p*-values denote results of a K-S test to compare drug-treatment data to data generated without drug treatment. (A) Mean number of generations in the no drug case. When fitting our model to data generated without drug treatment, we estimate a median of 4.24 and a mean of 4.75 generations. (B) Mean number of generations under treatment of 2’-C-meA. Fitting our model to data generated under the drug treatment of 2’-C-meA, we estimate a median of 4.67, and a mean of 5.06 generations. (C) Mean number of generations under treatment of rupintrivir. Fitting our model to data generated under the drug treatment of rupintrivir, we estimate a median of 5.77 and a mean of 6.0 of generations. (D) Mean number of generations under treatment of ganetespib. Fitting our model to data generated under the drug treatment of ganetespib, we estimate a median of 4, and a mean of 4.5 generations.

## Notes

### Competing Interest Statement

The authors have declared no competing interest.

